# Validation of an antigenic site targeted by monoclonal antibodies against Puumala virus

**DOI:** 10.1101/2023.08.10.552746

**Authors:** Alexander Plyusnin, Ashwini Kedari, Ilona Rissanen, Rommel Paneth Iheozor-Ejiofor, Åke Lundkvist, Olli Vapalahti, Lev Levanov

**Affiliations:** University of Helsinki, Department of Virology, Medicum, Helsinki, Finland; Institute of Biotechnology, Helsinki Institute of Life Science HiLIFE, University of Helsinki, Finland; Department of Medical Biochemistry and Microbiology, Microbiology-Immunology, Uppsala University, Sweden; Department of Virology and Immunology, HUSLAB, Helsinki University Hospital, Helsinki, Finland; Department of Veterinary Biosciences, University of Helsinki, Helsinki, Finland

**Keywords:** Monoclonal antibody, neutralizing epitope, site-directed mutagenesis

## Abstract

Identification of B-cell epitopes facilitates the development of vaccines, therapeutic antibodies and diagnostic tools. Previously, the binding site of the bank vole monoclonal antibody (mAb) 4G2 against Puumala virus (PUUV, an orthohantavirus in the *Hantaviridae* family of the *Bunyavirales* order) was predicted using a combination of methods, including pepscan, phage-display, and site-directed mutagenesis of vesicular stomatitis virus (VSV) particles pseudotyped with Gn and Gc glycoproteins from PUUV. These techniques led to the identification of the neutralization escape mutation F915A. To our surprise, a recent crystal structure of PUUV Gc in complex with Fab 4G2 revealed that residue F915 is distal from epitope of mAb 4G2. To clarify this issue and explore potential explanations for the inconsistency, we designed a mutagenesis experiment to probe the 4G2 epitope, with three PUUV pseudoviruses carrying amino acid changes E725A, S944F, and S946F, located within the structure-based 4G2 epitope in the Gc. These amino acid changes were able to convey neutralization escape from 4G2, and S944F and S946F also conveyed escape from neutralization by human mAb 1C9. Furthemore, our mapping of all the known neutralization evasion sites from hantaviral Gcs onto PUUV Gc revealed that over 60% of these sites reside within or close to the epitope of mAb 4G2, indicating that this region represents a crucial area targeted by neutralizing antibodies against various hantaviruses. The identification of this site of vulnerability could guide the creation of subunit vaccines against PUUV and other hantaviruses in the future.

## Full Text

Antibodies neutralize viruses via binding to viral proteins at sites referred to as B-cell epitopes, which can be categorized into two groups: linear and conformational. Linear epitopes consist of sequential continuous amino acid residues, whereas conformational epitopes consist of residues that are discontinuous in the protein sequence yet come within close proximity to form an antigenic surface on the three-dimensional structure of the protein. Identification of the exact location of B-cell epitopes facilitates the development of vaccines and selecting high-affinity antibodies for immunotherapy and immunodiagnostics (1, 2, 3). Most of the existing experimental methods for epitope mapping are laborious and time consuming, and fail to identify all epitopes. To verify mapping based on peptide scanning or mutagenesis analysis, further epitope validation by X-ray crystallography of cryogenic electron microscopy (cryoEM) may be required (4). This study explores the epitope of a potently neutralizing monoclonal antibody (mAb) 4G2 against Puumala virus (PUUV), the causative agent of *nephropathia epidemica*, a mild form of hemorrhagic fever with renal syndrome, across Northern Europe and Russia (5). Previously several putative mAb 4G2 binding sites were mapped on the surface of the PUUV Gc glycoprotein using pepscan and phage display techniques (6) and site-directed mutagenesis of antibody-reactive site _904_KCAFATTPVCQFDGNTIS_921_ (7), where the replacement of a conserved phenylalanine with alanine (F915A) resulted in a neutralization escape phenotype in vesicular stomatitis viruses (VSV) pseudotyped with PUUV Gn and Gc (7). These findings supported the idea that this region might contribute to the epitope of mAb 4G2.

Recently, however, Rissanen et al., determined the crystal structure of the PUUV Gc ectodomain in complex with the antigen-binding fragment (Fab) of 4G2 to 3.50 Å resolution (8). They showed that the antibody recognizes PUUV Gc at the junction of domains I and II of the type II fusion glycoprotein, distal from residue 915 (Figure 1A). It was demonstrated that R100 of complementarity determining region 3 (CDR3) of Fab 4G2 heavy chain is a key contributor to the interface that forms multiple hydrogen bonds with proximal Gc residues, including E725, I726 and S946. Notably, most of the residues comprising the epitope are buried in the post-fusion conformation of the Gc, leading the authors to propose a model whereby Fab 4G2 targets pre-fusion Gc and neutralizes through steric preclusion of fusogenic conformational states. In this work we applied site-directed mutagenesis to the epitope revealed by the crystal structure. The properties of mutant PUUV-VSV-pseudoviruses were analyzed in a neutralization assay, and the data obtained in the current study and in our previous work were assessed and compared with those obtained by Rissanen and colleagues.

**Figure 1.**
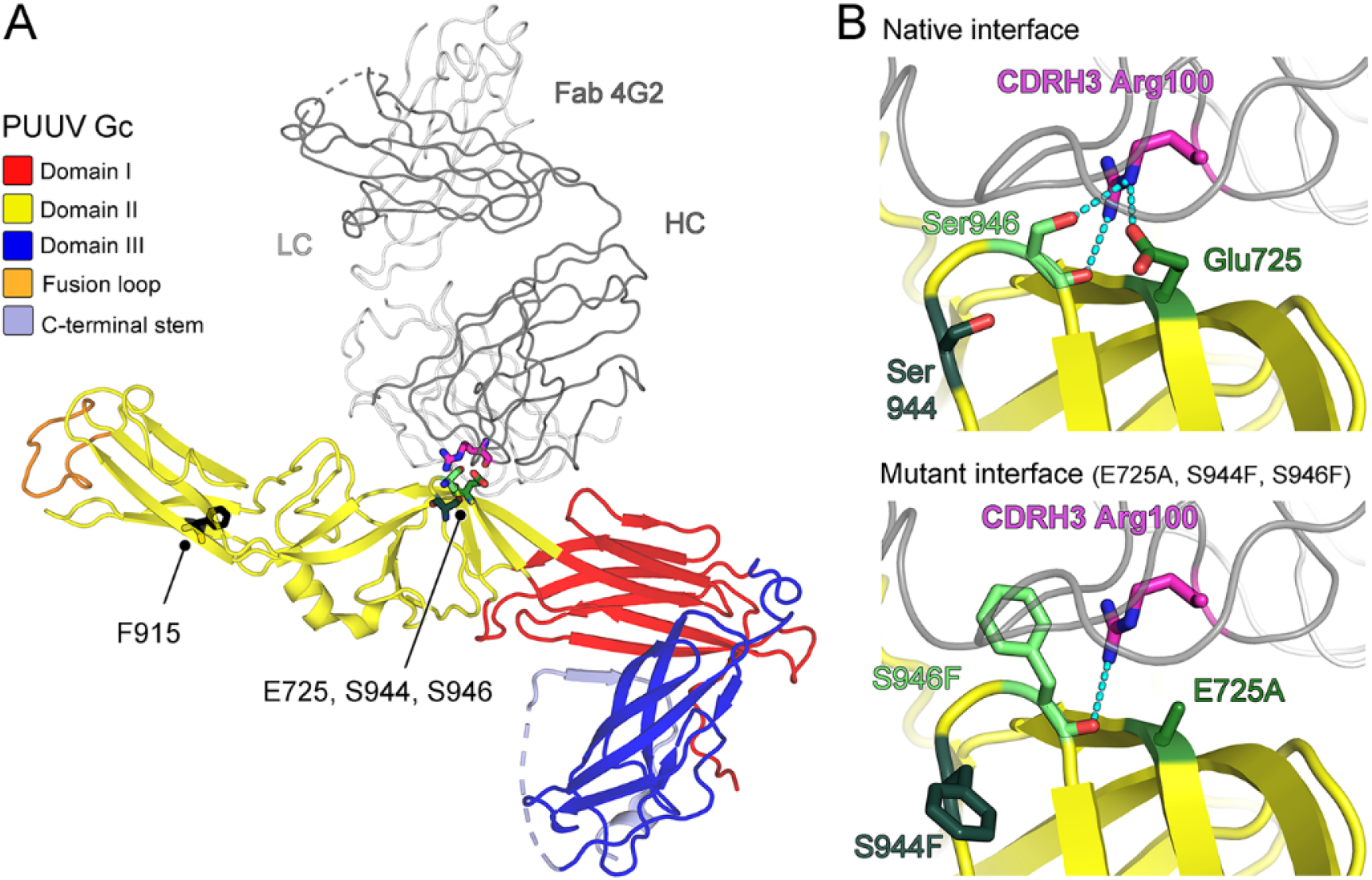
The complex of Fab 4G2 and PUUV Gc glycoprotein (PDB 6Z06) shows the structural context of neutralization escape mutation sites. A) Ribbon representation of PUUV Gc-Fab 4G2 complex, colored red, yellow, and blue for domains I-III, respectively. Amino acid F915 (black), previously identified as a neutralization escape site against mAb 4G2 (7), is highlighted alongside the three amino acids mutated in this study (E725, S944 and S946) and their interaction partner R100 (magenta) of CDR3 of heavy chain of Fab 4G2. B) A close-up view of the Fab-Gc interface, focused on E725, S944 and S946 and their hydrogen bond interactions with Fab 4G2 R100 (magenta). While not forming hydrogen bonds with the antibody, S944 most likely stabilizes epitope. Both native residues (upper panel) and the amino acids changes introduced in this study (lower panel) are presented.

Here, our mutagenesis experiments targeted three residues (Figure 1B): glutamic acid E725 and serine S946, which form hydrogen bonds with Fab 4G2 heavy chain R100 in the context of the crystal structure, and serine S944, since a previous study has shown that substitution S944F results in escape from neutralization by the human anti-PUUV mAb 1C9 (9). To test the impact of these residues on mAb 4G2 binding, E725 was replaced with an alanine, and S944 and S946 were replaced with phenylalanines. The mutations were introduced to plasmid pS7b (10) encoding Gn and Gc glycoproteins of PUUV using Phusion site-directed mutagenesis kit (Thermo Fisher Scientific), and VSV was pseudotyped with the mutant glycoproteins as described previously (7, 10). The resulting PUUV-VSV pseudotypes were able to infect Vero E6 cells (Figure 2). In order to evaluate the impact of amino acid changes E725A, S944F, and S946F on the neutralization characteristics of mAb 4G2, pseudotyped viruses were tested in a neutralization assay, as previously established (7, 10, 11), in the presence of mAb 4G2. Briefly, mAb was serially diluted four-fold (starting from 0.2 mg/ml) and mixed with an equal volume containing 150 fluorescence units of VSV pseudotypes. The mixture was incubated for 1 hour at 37 °C and then administered onto Vero E6 cell monolayers in 96-well tissue culture plates. Following a one-hour adsorption period, the inoculum was exchanged with Eagle’s minimum essential medium. After 18 hours, cells that had been infected with fluorescent PUUV-VSV pseudotypes were examined and counted. In-line with their putative role at the antibody interface, amino acid changes E725A, S944F, and S946F each resulted in mAb 4G2 escape phenotypes in neutralization assay (Figure 3A). These data confirm the functional relevance of this region for mAb 4G2 recognition. The amino acid change F915A, observed to convey neutralization escape in our previous study (7), likely impacted the folding of PUUV Gc to an extent where the epitope of mAb 4G2 was affected, despite the remoteness of residue 915 from the epitope. It should also be noted that most epitope candidate peptides found by pepscan and phage display for mAb 4G2 in a previous study (6) do not fall within the epitope determined by crystallography (8).

**Figure 2.**
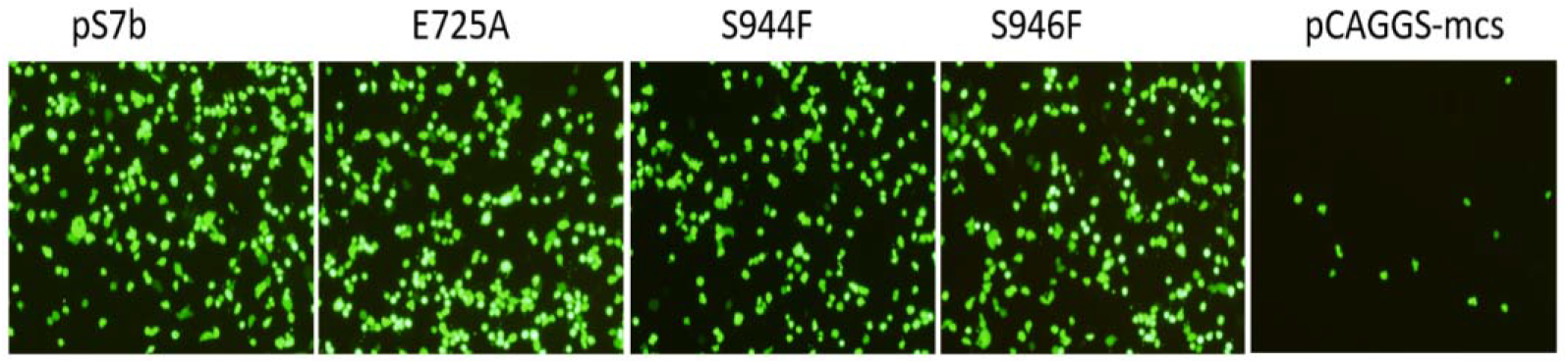
Expression of EGFP in Vero E6 cells infected with VSV _Δ_G EGFP particles pseudotyped with native (pS7b) and mutant (E725A, S944F, S946F) PUUV glycoproteins. The empty plasmid pCAGGS-mcs was used as a negative control in pseudotyping experiment. Infection was done with 50μ of diluted (10^−1^) virus stock in 96-well plate. After a 1 hour incubation period, the inoculum was removed, and fresh Eagle’s minimum essential medium was added, and the cells were incubated at 37ºC in a CO_2_ incubator. At 24 h postinfection, the GFP-expressing cells were examined under a fluorescence microscope

**Figure 3.**
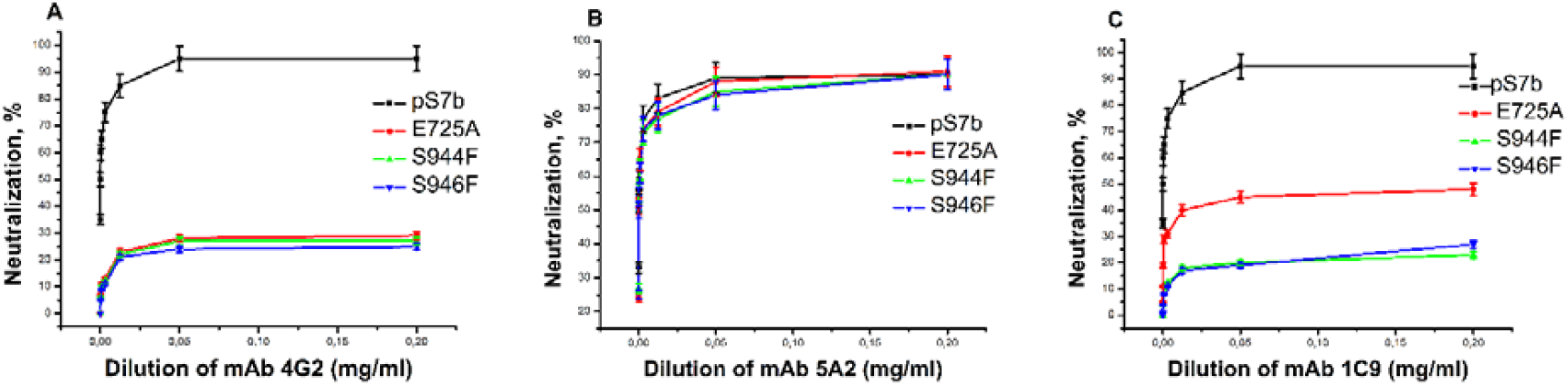
Neutralizing activity of mAb 4G2 (A), 5A2 (B) and 1C9 (C) against PUUV with VSV particles pseudotyped with native (ps7b) and mutant (E275A, S944F, S946F) PUUV glycoproteins. mAbs were serially diluted four-fold (starting from 0.2 mg/ml) and mixed with an equal volume containing 150 fluorescence units of VSV pseudotypes. A total of 50□μl of medium containing 150 fluorescence units of VSV pseudotypes was incubated with an equal volume of serially diluted mAbs for 1□h at 37□°C. Then 90□μl of the mixture was inoculated onto Vero E6 cells monolayers in 96-well tissue culture plates. After adsorption for 1 h, the inoculum was replaced with Eagle’s minimum essential medium. After 18□h, cells infected with fluorescent VSV pseudotypes were examined and counted. Each experiment was done in triplicate. Titres are presented as mean values ± standard deviations. P□0.05. Statistical analysis was conducted using one-way ANOVA implemented in Origin software.

To confirm the antibody specificity of the epitope, the E725A, S944F and S946F pseudotypes were also tested in a neutralization assay with bank vole mAb 5A2 (6, 7) that targets the Gn glycoprotein of PUUV and human mAb 1C9 (6, 7) against the Gc glycoprotein of PUUV. As expected, mAb 5A2 effectively inhibited infection by all the mutant PUUV-VSV pseudotypes (Figure 3B). In contrast, mutant pseudotypes S944F and S946F escaped from neutralization by mAb 1C9 (Figure 3C), indicating that bank vole mAb 4G2 and human mAb 1C9 share a similar B-cell epitope on the Gc glycoprotein of PUUV. We also observed that substitution E725A conveyed partial escape from mAb 1C9 (Figure 3C), suggesting that this residue is involved in 1C9 binding, but to a lesser extent than in mAb 4G2 binding.

To understand whether this region, implicated in antibody recognition for both bank vole and human anti-PUUV mAbs, is widely targeted across hantaviruses, we mapped all known neutralization escape sites discovered in hantaviral Gc proteins onto the Gc protein of PUUV. While the crystal structure of PUUV Gc-Fab 4G2 (PDB 6Z06) remains the only structure of a neutralizing antibody-hantaviral Gc published to date, recent studies have greatly expanded the portfolio of neutralizing antibodies targeted against hantaviral Gn and Gc proteins and their associated neutralization escape sites (12, 13, 14). Our literature search identified 24 amino acid changes in the Gc that convey neutralization escape from antibodies against PUUV, Hantaan virus (HNTV), Andes virus (ANDV), and Sin Nombre virus (SNV) (Supplementary Table 1). We note that 21 of the 24 sites were reported in 2020 or later (12, 13, 14), allowing us to perform the most comprehensive mapping of Gc neutralization escape mutation sites to date. In order to examine these sites in the context of the hantaviral surface, which carries a lattice of hetero-octameric hantaviral spikes, an individual (Gn/Gc)_4_ spike was generated from on the published model of PUUV Gc and PUUV Gn fitted into a cryoEM reconstruction of the hantaviral surface (PDB7B0A, EMD-11966) (8) in UCSF ChimeraX (15) by applying C4 symmetry. Homologous positions in PUUV Gc were determined for neutralization escape sites identified in the Gc of HTNV, ANDV or SNV, and all sites were mapped onto the PUUV spike in PyMOL (The PyMOL Molecular Graphics System, Version 2.5.0, Schrödinger, LLC), along with the structure-based epitope of mAb 4G2 (Figure 4). In line with its expected accessibility to antibodies, escape sites are accumulated on the membrane-distal face of the Gc. Our mapping reveals that the hinge region between Gc domains I and II, targeted by mAb 4G2, is a hotspot for antibody binding, and carries 15 of the 24 neutralization escape sites (62.5%). Half of all sites (12/24) are located within or directly adjacent to the mAb 4G2 epitope, which itself comprises only ∼4% (830 Å^2^ of 21242 Å^2^) of the total surface of Gc. The sites concentrated in the mAb 4G2 epitope region convey escape from antibodies against PUUV, HTNV, ANDV and SNV, indicating that in addition to its apparent role as a major antigenic epitope, this region may comprise a site of vulnearability to neutralization across orthohantaviruses. In conclusion, we applied site-directed mutagenesis to probe key residues at the conformational epitope of mAb 4G2 determined by crystallography (6). Amino acid changes E725A, S944F and S946F were able to convey neutralization escape to PUUV-VSV pseudotypes, and confirm the functional relevance of this region for bank vole mAb 4G2 binding. Furthermore, S944F and S946F amino acid changes convey neutralization escape from human mAb 1C9, and a comprehensive mapping of neutralization escape sites found in hantaviral Gc proteins revealed that the region targeted by mAb 4G2 is likely to play a major role in neutralizing antibody recognition across hantaviral species.

**Figure 4.**
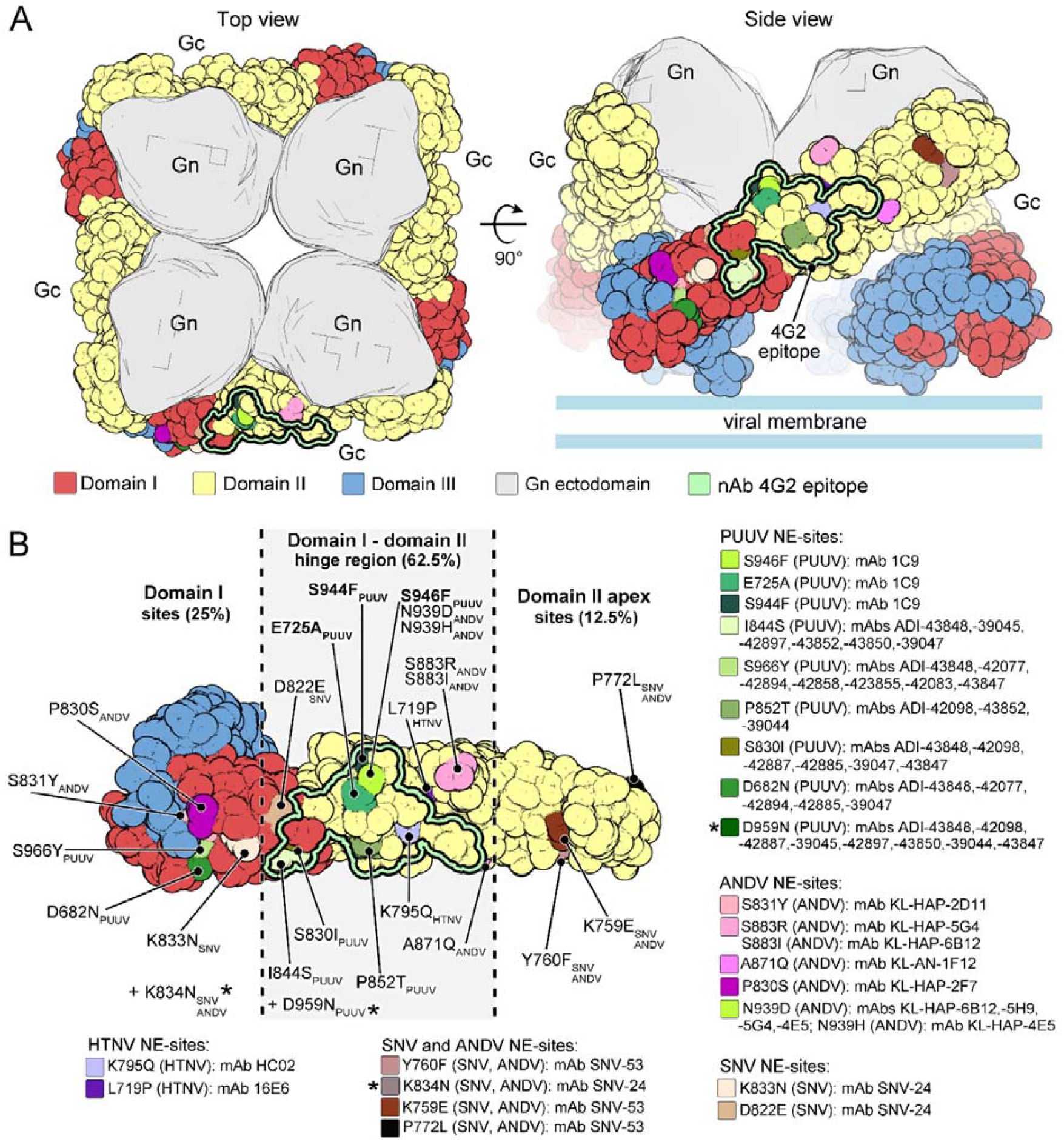
The distribution of neutralization escape mutation sites in the Gc indicates a key antigenic site. A) Representation of the PUUV (Gn/Gc)_4_ spike with the Gn shown as grey density and Gc shown as solvent-accessible surface, coloured red, yellow, and blue for domains I-III, respectively. Reported neutralization escape (NE) mutation sites (Supplementary Table 1) and the epitope of neutralizing antibody (nAb) 4G2 are shown mapped onto the surface of a single PUUV Gc. B) Detailed view of the NE mutation sites for antibodies against PUUV, HNTV, ANDV, and SNV on the PUUV Gc single ectodomain. The nAb 4G2 epitope is delineated by a light green borderline. PUUV, ANDV, SNV, and HTNV NE sites are shown in shades of green, pink, beige, and purple respectively. Shared ANDV and SNV NE sites are illustrated in shades of brown. Two NE sites denoted as *K834N and *D959N are positioned on the membrane –proximal side of the Gc, and therefore, are not visible on the shown solvent-facing view of the spike. Site F915A was not included in the mapping, as the crystal structure of the PUUV Gc-Fab 4G2 complex has revealed this site to not be indicative of an antibody binding region.

## Acknowledgements

We thank Dr John Rose (Yale University) for VSV rescue plasmids. We appreciate the excellent technical assistance of Irina Suomalainen.

## Conflicts of interest

The authors declare that there are no conflicts of interest.

### Abbreviations

mAb: monoclonal antibody
PUUV: puumala virus
VSV: vesicular stomatitis virus
EGFP: enhanced green fluorescent protein.

