## Supplemental Table 1 for "Validation of an antigenic site targeted by monoclonal antibodies against Puumala virus"

**Supplementary Table 1**. Overview of the mAb 4G2 epitope and neutralization escape (NE) mutation sites identified on hantaviral Gc proteins.

| Site location | Virus | Epitope or NE site | Associated antibody | Reference |
|---|---|---|---|---|
| **Structure-based mAb 4G2 epitope** | | | | |
| Gc domain I and II | PUUV | Residues 723-727, 805-810, 829-831, 844-845, 851-854, 870-874, 876, 944-947 | mAb 4G2 epitope; PDB 6Z06 | Rissanen, I., et al. Elife, 2020. **9**:e58242. |
| **NE-sites reported before 2020** | | | | |
| Gc domain II | HTNV | L719P | mAb 16E6 | Wang, M., et al. Virology, 1993. **197**(2): p. 757-66. |
| Gc domain II | HNTV | K795Q | mAb HC02 | Wang, M., et al. Virology, 1993. **197**(2): p. 757-66. |
| Gc domain II | PUUV | S944F | mAb 1C9 | Hörling, J. and A. Lundkvist, Virus Res, 1997. **48**(1): p. 89-100. |
| **NE-sites reported 2020 onwards** | | | | |
| Gc domain I and II | SNV<br>ANDV | D822E, K833N, K834N<br>K834N, K759E/P772L<br>Y760F, K759E/P772L | mAb SNV-24<br>mAb ANDV-44<br>mAb SNV-53 | Engdahl, T.B., et al. Elife, 2023. **12**:e81743. |
| Gc domain I and II | PUUV | D682N, S830I, I844S, D959N, S966Y<br>D682N, S966Y<br>D682N<br>S830I, P852T, D959N<br>S830I, I844S, D959N<br>I844S, D959N<br>I844S, P852T<br>P852T, D959N<br>D959N<br>D682N, S830I, I844S, D959N, S966Y<br>S830I | mAb ADI-43848<br>ADI-42077, ADI-42894, ADI-42894<br>ADI-42885<br>ADI-42098<br>ADI-42887<br>ADI-39045, ADI-42897<br>ADI-43852<br>ADI-43850<br>ADI-39044, ADI-42858, ADI-42855<br>ADI-39047<br>ADI-43847 | Mittler, E., et al. Sci Transl Med, 2022. **14**(636): p. eabl5399. |
| Gc domain I and II | ANDV | P830S<br>S831Y<br>S883I<br>S883R<br>A871Q<br>N939D<br>N939H | mAb KL-HAP-2F7<br>KL-HAP-2D11<br>KL-HAP-6B12<br>KL-HAP-5G4<br>KL-AN-1F12<br>KL-HAP-6B12, KL-HAP-5H9, KL-HAP-5G4<br>mAb KL-HAP-4E5 | Duehr, J., et al. mBio, 2020. **11**(2). |
